# Sex-specific role for the long noncoding RNA *Pnky* in mouse behavior

**DOI:** 10.1101/2023.12.05.569777

**Authors:** Parna Saha, Rebecca E. Andersen, Sung Jun Hong, Eugene Gil, Jeffrey Simms, Daniel A. Lim

**Affiliations:** Department of Neurological Surgery; Eli and Edythe Broad Center of Regeneration Medicine and Stem Cell Research; Behavioral Core, Gladstone Institutes, San Francisco; San Francisco Veterans Affairs Medical Center; University of California, San Francisco, San Francisco, CA 94143, USA

## Abstract

The human brain expresses thousands of different long noncoding RNAs (lncRNAs), and aberrant expression of specific lncRNAs has been associated with cognitive and psychiatric disorders. While a growing number of lncRNAs are now known to regulate neural cell development and function, relatively few have been shown to underlie animal behavior, particularly with genetic strategies that establish lncRNA function in *trans*. *Pnky* is an evolutionarily conserved, neural lncRNA that regulates brain development. Using mouse genetic strategies, we show that *Pnky* has sex-specific roles in mouse behavior and that this lncRNA underlies specific behavior by functioning in *trans*. Male *Pnky*-knockout (KO) mice have deficits in cued fear recall, a type of Pavlovian associative memory. In female *Pnky*-KO mice, the acoustic startle response (ASR) is increased and accompanied by a decrease in prepulse inhibition (PPI), both of which are behaviors altered in affective disorders. Remarkably, expression of *Pnky* from a bacterial artificial chromosome (BAC) transgene reverses the ASR phenotype of female *Pnky*-KO mice, demonstrating that *Pnky* underlies specific animal behavior by functioning in *trans*. More broadly, these data provide genetic evidence that a lncRNA gene and its function in *trans* can play a key role in the behavior of adult mammals, contributing fundamental knowledge to our growing understanding of the association between specific lncRNAs and disorders of cognition and mood.

## INTRODUCTION

Long noncoding RNAs (lncRNAs) –– transcripts longer than 200 nucleotides (nts) that do not encode proteins –– have emerged as key regulators of important biological processes (Rinn and Chang 2012; Statello et al. 2021). Of the tens of thousands of distinct lncRNAs produced by the mammalian genome, many are brain and neural cell type specific (Andersen and Lim 2018; Mercer et al. 2008; Liu et al. 2016; Molyneaux et al. 2016; Ramos et al. 2013), and aberrant lncRNA expression has been implicated in a broad range of neurological disorders including schizophrenia, depression, autism, and Alzheimer’s disease (S. Yang et al. 2021; Parikshak et al. 2016). However, although the list of lncRNAs experimentally shown to regulate neural cell biology is expanding (Ang, Trevino, and Chang 2020; Aliperti, Skonieczna, and Cerase 2021; Wu et al. 2022), very few lncRNAs have been demonstrated to have roles in animal behavior (Ip et al. 2016; Issler et al. 2020; Kukharsky et al. 2020; Labonté et al. 2021; Issler et al. 2022; Rontani et al. 2021; Barry et al. 2014; Keihani et al. 2019).

*Pnky* is an evolutionarily conserved, neural lncRNA that regulates mouse neural stem cell (NSC) function both *in vitro* and *in vivo* (Ramos et al. 2015; Andersen et al. 2019). *Pnky* conditional knockout (cKO) (Andersen et al. 2019) or transcript knockdown (KD) (Ramos et al. 2015; Lin et al. 2014) increases neuronal production from cultured mouse NSCs, and other studies indicate a role for *Pnky* in NSC migration (Du et al. 2023). *In vivo*, genetic deletion of *Pnky* – either cKO or germline KO – causes aberrant cortical development, resulting in alterations of neuron subtype abundance (Andersen et al. 2019).

Whether *Pnky*-KO would have an effect upon behavior was very unclear. Only a small number of lncRNAs have been genetically disrupted in mice, and in certain well-studied cases, behavior is only mildly affected (Ip et al. 2016) or apparently not at all (Oliver et al. 2015).

Although the cellular phenotype of acute *Pnky*-deletion in cultured NSCs is dramatic (*e.g.,* ∼4-fold increase in neuronal differentiation), *Pnky*-deletion *in vivo* produces relatively subtle changes in cortical anatomy (*e.g.,* ∼15% difference in specific layers of the cortex), suggesting that the *in vivo* context provides compensatory molecular mechanisms for the absence of *Pnky* in neurodevelopment (Ramos et al. 2015; Andersen et al. 2019). Given that animal behavior includes many additional layers of potential compensatory mechanisms (*e.g.,* the plasticity of neuronal circuits), observing a behavioral phenotype in a neural lncRNA-KO mouse is a relatively demanding benchmark to establish the importance of that lncRNA.

Gene-disrupting manipulations to the genome can disrupt potential *cis* function of the locus. For the study of lncRNA function, this consideration is particularly important since many lncRNAs appear to function in *cis*, regulating the expression of neighboring genes (Bassett et al. 2014; Engreitz et al. 2016). Even the use of KD approaches (*e.g*., short-hairpin RNAs and antisense oligonucleotides) does not distinguish *cis* from *trans* function, since transcript KD can disrupt transcriptional elongation, which itself modulates neighboring gene expression (Kopp and Mendell 2018). In previous studies, neither *Pnky*-deletion nor *Pnky* KD alters the expression level of any other gene within a 1 MB chromosome window, suggesting that *Pnky* does not function in *cis* (Ramos et al. 2015; Andersen et al. 2019)

LncRNA expression from a BAC transgene can be used to test for *trans* function and further demonstrate function of the lncRNA itself (Bassett et al. 2014; Kopp and Mendell 2018). For example, genetic disruption of lncRNA *Fendrr* causes defects in heart development, and expression of *Fendrr* from a BAC transgene rescues some of the developmental abnormalities, indicating that *Fendrr* can function in *trans* (Grote et al. 2013). In a previous study (Andersen et al. 2019), we generated a transgenic mouse line that expresses *Pnky* from a ∼170 kb BAC insert (BAC-*Pnky*) that lacks other known genes, and BAC-*Pnky* rescues transcriptomic, cell culture, and *in vivo* developmental phenotypes of *Pnky*-deletion. These results strongly support the notion that *Pnky* encodes a *trans*-acting functional lncRNA. However, whether the *trans*-action of *Pnky* can also underlie the much more complex phenotypes of animal behavior was unknown, and *trans* rescue of a lncRNA-KO behavioral phenotype remains a highly stringent benchmark – one that to the best of our knowledge has not yet been achieved – to demonstrate lncRNA function.

In this study, we investigated the role of *Pnky* in adult mouse behavior with *Pnky*-KO and BAC-*Pnky* mice. We discovered sex-specific phenotypes of *Pnky*-KO in animal behaviors that relate to cognition and certain affective disorders. Furthermore, we found that BAC-*Pnky* can selectively reverse an important *Pnky*-KO behavioral phenotype, also in a sex-specific manner. In addition to characterizing *Pnky* as a lncRNA important to animal behavior, these data more broadly illustrate how lncRNAs can underlie cognitive and psychiatric disorders.

## RESULTS

### *Pnky*-KO does not disrupt basic neurological function and nesting behavior

*Pnky*-KO mice are born at expected Mendelian frequencies and do not have obvious major deficits of general health and neurological behavior (Andersen et al. 2019). To more systematically assess behavior, we analyzed a cohort of *Pnky*-KO (n= 12 females and 14 males) mice and littermate *Pnky+/+* (WT, n= 11 females and 11 males) controls with a panel of behavioral tests 5-6 months after their birth (**Figure 1A**). Data from behavioral tests were collected by experimenters blinded to the genotype.

**Figure 1:**
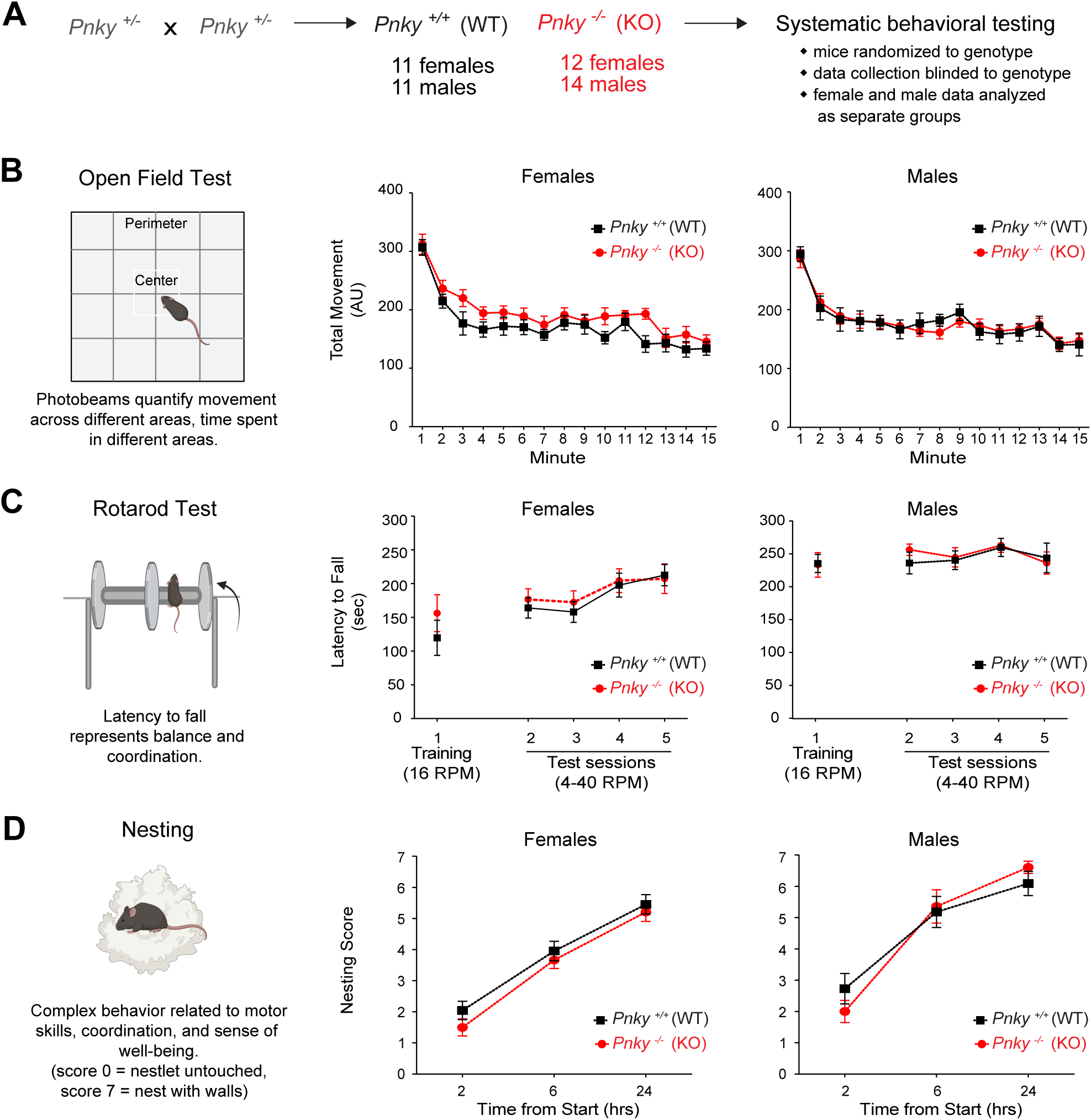
*Pnky*-KO does not result in basic neurological deficits. **A**) *Pnky*+/-mice were crossed to obtain a cohort of *Pnky*-KO (n= 12 females and 14 males) mice and littermate *Pnky+/+* (WT, n= 11 females and 11 males). **B)** Open field test assesses the gross activity levels (locomotion) and natural exploration habits in rodents. Total movements (measured in arbitrary units (AU)) in the open field (center and outer zones) are comparable between *Pnky*-WT and *Pnky-* KO animals (repeated measures two-way ANOVAs, *p =* ns). **C)** Rotarod test measures balance and motor coordination. Latency to fall on the Rotarod test in the training (Welch’s *t*-tests, *p* = ns) and test phases (repeated measures two-way ANOVAs, *p =* ns) shows no difference between the two genotypes. Rotarod mean session data is reported, each session is comprised of 3 trials separated by a 15-20 min ITI. **D)** Nest building is an innate behavior to assess general well-being. Nesting behavior is unaffected by *Pnky* deletion across the 24 hr testing period (repeated measures two-way ANOVAs, *p =* ns). ns = non-significant, data is represented as mean ± SEM.

To assess locomotor function, we performed the open field test (OFT) (Kraeuter, Guest, and Sarnyai 2019) in which ambulation within and across different zones in an open, wall-enclosed area is measured using photobeam arrays. The amount of total movement, ambulatory movement and rearing behavior was not different between *Pnky*-KO mice and controls (**Figure 1B, Figure1 Supplement 1 A-B**), indicating normal levels of spontaneous motor activity across the different experimental groups.

To evaluate motor coordination and balance, we employed the rotarod test (Shiotsuki et al. 2010). Neither male nor female *Pnky*-KO mice exhibited differences in their latency to fall in the training (constant speed, 16 rotations per minute (RPM)) or the testing (accelerating, 4-40RPM) phases as compared to their WT controls (**Figure 1C**).

Nest building is an important task for rodents that requires complex motor skills and spatial memory. Impaired nest building is also a sensitive indicator of decreased general health and sense of well-being (Jirkof 2014). In both male and female mouse cohorts, the nesting score of *Pnky*-KO mice was not different from that of their WT littermates (**Figure 1D**). Thus, *Pnky* is not required for general locomotor function, motor coordination, balance, and the complex behavior of nest building.

### *Pnky*-KO mice do not exhibit changes in anxiety or deficits in social interactions

In OFT, the fraction of time spent in the open center of the field versus that of the walled perimeter is a measure of anxiety, and there was no difference with *Pnky*-KO (**Figure 2** **Supplement 1A**). To further investigate anxiety related behaviors, we used the elevated plus maze (EPM) test (Walf and Frye 2007). The EPM is a raised platform in the shape of a plus sign (+), consisting of two open arms (no walls) that are intersected by two enclosed arms (with walls). Mice with increased anxiety avoid the elevated, open spaces of the open arms and spend more time in the enclosed arms of the EPM. *Pnky*-KO mice were not different from controls in EPM testing (**Figure 2A**, **Figure 2** **Supplement 1B**), further indicating a normal, baseline level of anxiety in *Pnky*-KO mice.

**Figure 2.**
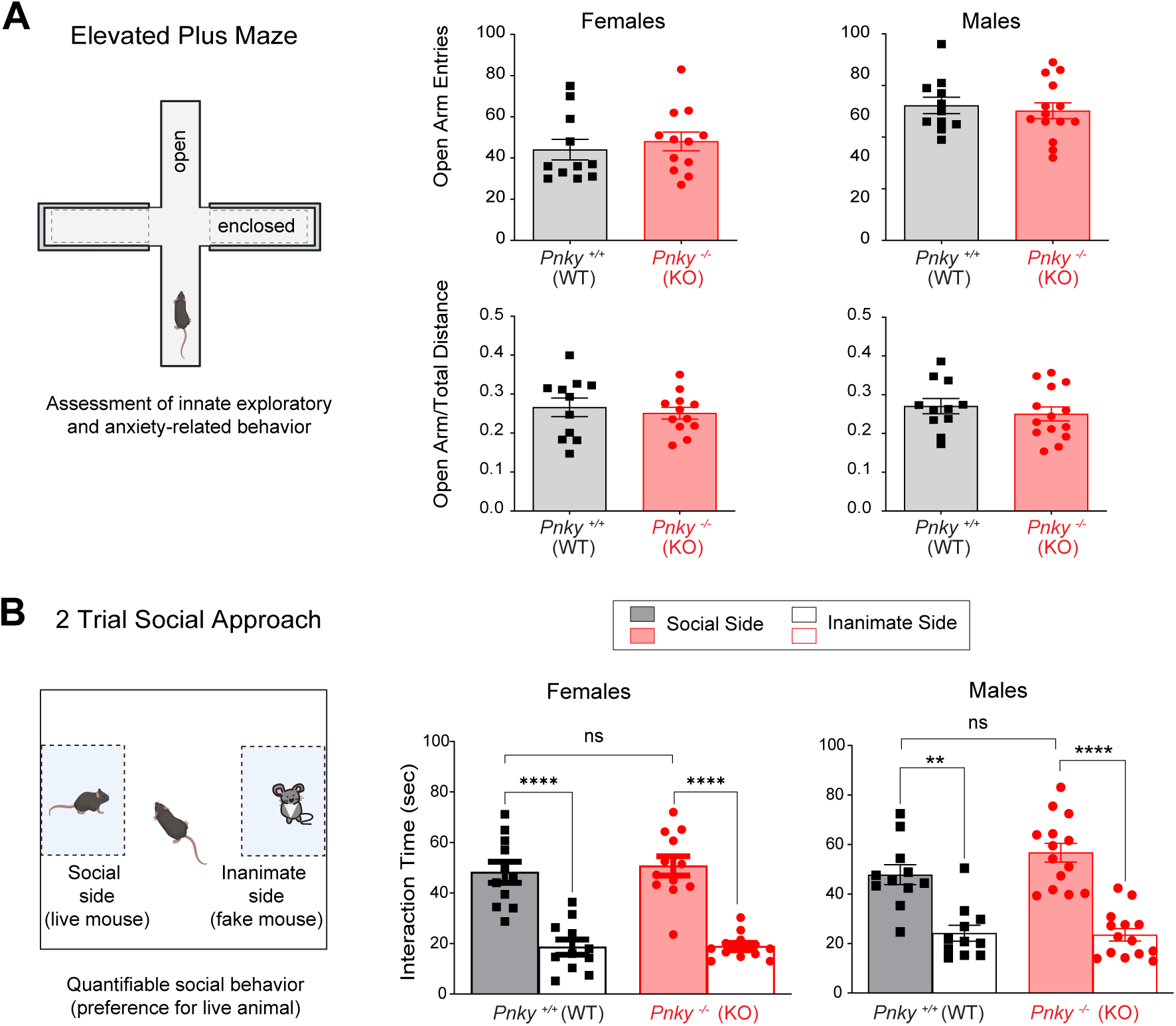
*Pnky*-KO mice do not exhibit increased anxiety or deficits in social interactions. **A**) The elevated plus maze (EPM) test is used to assess anxiety related behavior. In the EPM, the number of entries into the open arm (females, Mann-Whitney test, *p* = ns; males, Welch’s *t*-test, *p* = ns) and the ratio of open arm distance to total distance (Welch’s *t*-tests, *p* = ns) was equal for both the genotypes. **B)** The social approach test assesses general sociability and preference for social novelty. In the social approach trial, male and female mice of both genotypes show increased preference (interaction time in seconds) for the social side whereas the difference in interaction time across genotypes is non-significant. Females: Paired t test \*\*\*\**p* = 0.0001 for *Pnky*-WT and *Pnky*-KO females social vs inanimate; Pnky-WT vs *Pnky*-KO social interaction, Welch’s *t*-test, *p* = ns). Males – Paired t test \*\**p*= 0.0029 for social vs inanimate for *Pnky*-WT males; paired *t* test \*\*\*\**p*= 0.0001 social vs inanimate for *Pnky*-KO males; Pnky-WT vs *Pnky*-KO social interaction, Welch’s *t*-test, *p* = ns). \*\**p* < 0.01, \*\*\*\**p* <0.0001, ns = non-significant, data is represented as mean ± SEM.

The two-chamber social approach test measures the preference of mice for interactions with other mice (M. Yang, Silverman, and Crawley 2011). For this test, mice were allowed to explore an arena that has two chambers, one of which contains a wire enclosure with an inanimate mouse toy (inanimate chamber) and the other containing a wire enclosure with a live mouse (social chamber). Mice with social deficits generally have decreased interaction bouts and time with the stimulus mouse as compared to the inanimate mouse. Both sexes of the *Pnky*-KO and WT mice interacted significantly more with the social chamber, and there were no differences between the two genotypes (**Figure 2B**), demonstrating a normal level of social interaction in *Pnky*-KO mice.

## Male *Pnky*-KO mice have deficits in cued fear recall

To investigate cognitive behavior, we employed two different testing paradigms that evaluate different aspects of learning and memory. The object-context congruence test measures visual recognition of objects and their association with different environmental contexts (Leger et al. 2013). After training to associate specific objects (blocks vs. flasks) with different environmental contexts (white walls vs. checkered pattern walls), mice were tested for whether they can distinguish “congruent” object-context associations (e.g., block object with white wall context) from “incongruent” associations (e.g., block object with checkered pattern wall context). *Pnky*-KO and WT mice did not exhibit significant differences in either the training or test phases of the object-context congruence test (**Figure 3** **Supplement 1**).

**Figure 3.**
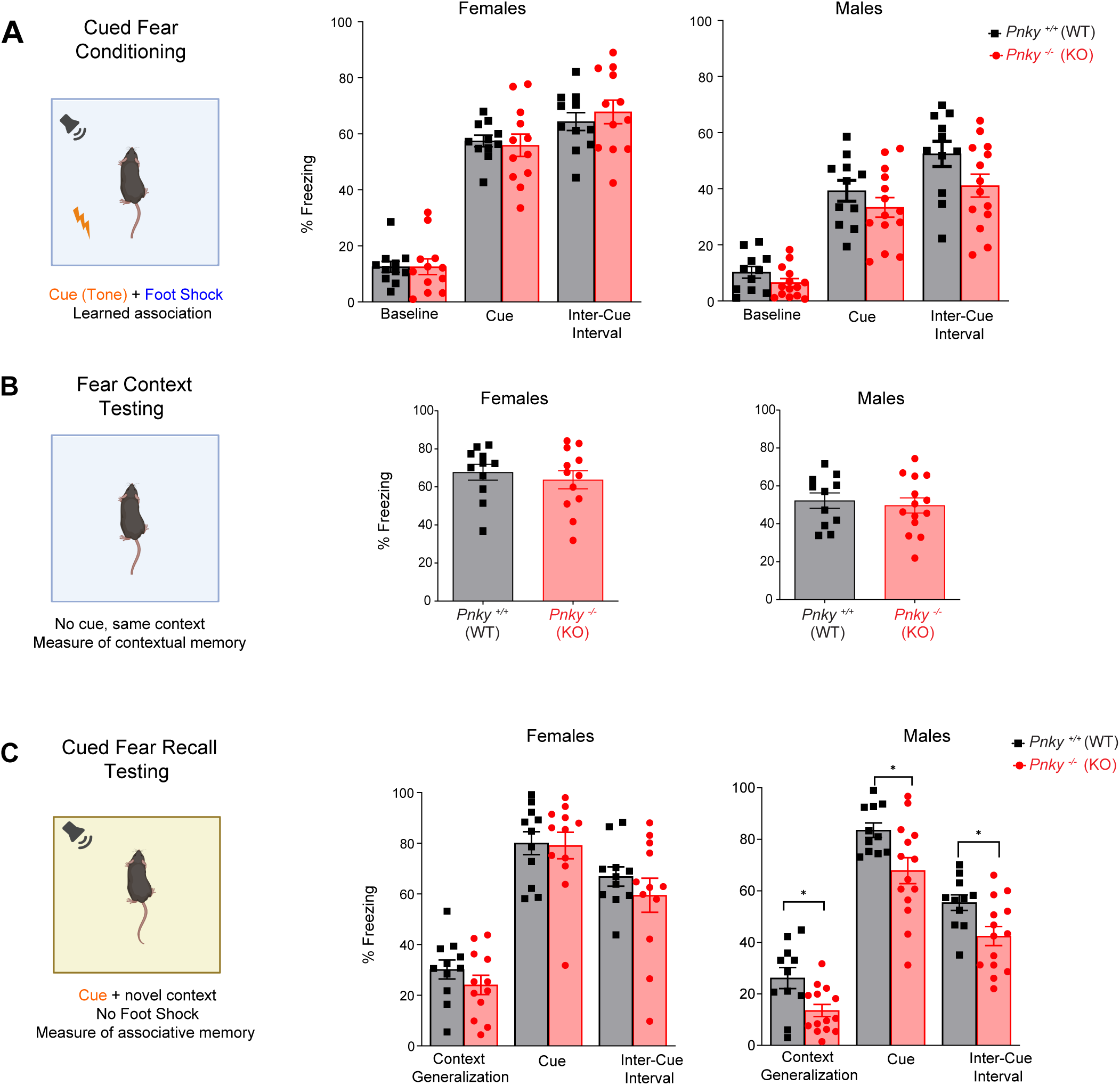
Male *Pnky*-KO mice have deficits in cued fear recall. **A**) Cued fear conditioning test assesses the ability of mice to learn and remember an association between environmental cues and aversive experiences measured by recording the freezing response. % Freezing (lack of movement aside from normal respiration) is calculated in 1 min blocks. **A)** During cued fear conditioning freezing behavior in *Pnky*-KO mice was comparable to its WT controls (multiple *t*-tests for the baseline, cue presentations, and ICI for female and male mice, *p* = ns). **B)** In the fear context test, the freezing behavior show no difference between between the *Pnky*-WT and *Pnky*-KO groups (Welch’s t-tests, *p =* ns). **C)** During cued fear recall test, *Pnky*-WT and *Pnky*-KO females exhibit no significant difference in freezing behavior (multiple *t*-tests for the generalization, cue presentations and ICI, *p* = ns). *Pnky-*KO male mice show decreased freezing compared to the WT control in the 5-minute generalization period multiple *t*-tests for the generalization (\**p* = 0.0101), cue presentations (\**p* = 0.0183), and ICI, (\**p* = 0.0152)). \**p* < 0.05, ns= non-significant, data is represented as mean ± SEM.

The cued fear conditioning and recall test measures the ability of mice to learn and remember an association between environmental cues and aversive experiences (Wehner and Radcliffe 2004). This test of Pavlovian conditioning consists of three phases: (1) cued fear training, (2) fear context testing and (3) cued fear recall testing. During the training phase, a tone is paired with a mild electric shock, and mouse freezing behavior is quantified by video analysis. In normal mice, with successive presentations of tone-shock pairs, the amount of freezing increases, which corresponds to associative learning. In both sexes, during cued fear training, the freezing behavior of *Pnky*-KO mice was not significantly different from their WT controls (**Figure 3A**, **Figure 3** **Supplement 2A**), indicating a similar degree of learning of this behavior among *Pnky*-KO and WT mice.

Fear context testing is performed 24 hours later in the same environmental context (same box as training phase) where freezing behavior is again quantified. This second phase evaluates whether mice have learned to associate the environmental context with the aversive experience (foot shock). To get a pure measure of contextual memory, no auditory cues or foot shocks are presented during this phase. During fear context testing, the amount of freezing behavior was not different between *Pnky*-KO and WT mice, regardless of sex (**Figure 3B**), indicating that *Pnky*-KO does not impair memory of the fear-context association.

The third phase – cued fear recall testing – occurs 24 hours after fear context testing. In this test, the visual and tactile appearance of the box is changed, which removes the prior environmental context associations. The same repetitive pattern of cues (tones) is presented but without any foot shocks, and mouse freezing behavior is quantified. In female *Pnky*-KO mice, the amount of freezing with the auditory cues was not different from that of WT controls (**Figure 3C**). In contrast, male *Pnky*-KO mice exhibited decreased freezing behavior in the 5-minute context generalization phase, during the tone presentations, and in the inter-cue intervals (ICI) (**Figure 3C**, **Figure 3** **Supplement 2B**), indicating a deficit of cued fear recall in male mice that lack *Pnky*. Thus, *Pnky* is required for a specific type of associated memory in a sex-specific manner.

### *Pnky*-KO increases the acoustic startle response and decreases prepulse inhibition in female mice

The acoustic startle response (ASR) in mice is a non-stereotypic behavior that is characterized by a quantifiable flinching behavior in response to a loud (100-120 dB) sound stimulus. Differences in baseline startle responses can occur with various psychiatric conditions and specific brain lesions (Angrilli et al. 1996; Poli and Angrilli 2015; Koch 1999). To measure ASR, mice are placed on a movement sensor plate (that measures the amplitude of the startle, flinching behavior) and exposed to a series of tones of varying intensities at random intervals for 15 minutes. Female *Pnky*-KO mice exhibited greatly increased startle amplitudes in response to tones across a broad range (80 to 120 dB) of acoustic intensities (**Figure 4A**). In male *Pnky*-KO mice, the startle amplitude at each tone intensity was not different from that of their WT controls (**Figure 4A**). Thus, loss of *Pnky* greatly increases the ASR of female mice.

**Figure 4.**
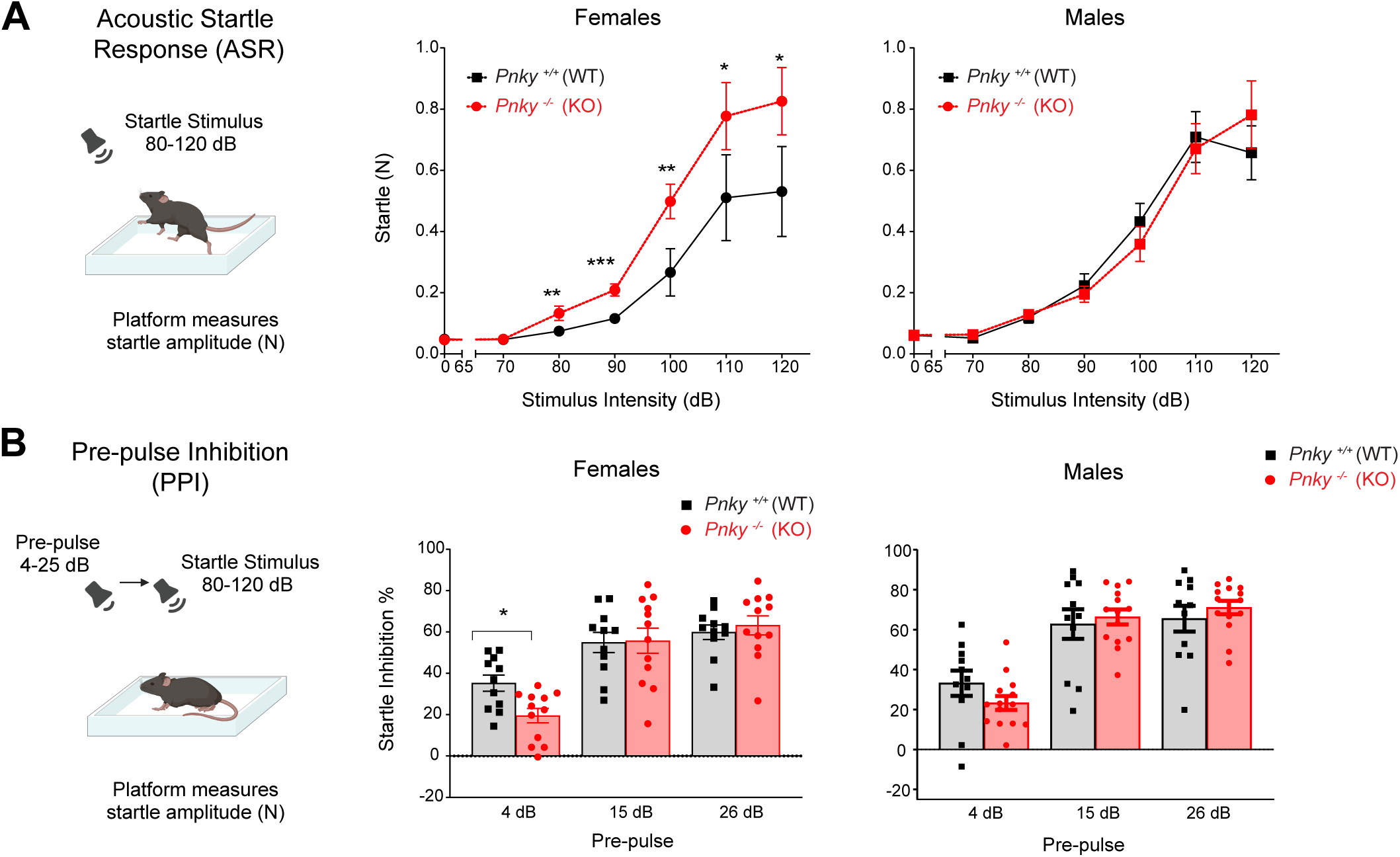
*Pnky*-KO increases the acoustic startle response and decreases prepulse inhibition in female mice. **A**) Acoustic startle response (ASR) measures quantifiable flinching behavior in response to a loud (80-120 dB) auditory stimulus. *Pnky* KO females exhibit increased startle with increasing intensity of the acoustic stimuli (rank summary analysis overall effect *p =* 0.0022 across the whole range of stimuli, followed by *t*-tests at 0dB – ns, 70dB – ns, 80dB – \*\**p* = 0.0045, 90 dB – \*\*\**p* = 0.0009, 100 dB – \*\**p* = 0.0045, 110 dB – *, *p* = 0.0439, 120dB – \**p* = 0.0439). Startle response in Pnky KO males was unchanged (rank summary analysis overall effect *p* = ns). **B)** Prepulse inhibition (PPI) of ASR is the reduction in startle response when a weak, non-startling tone (prepulse) precedes the auditory startle stimulus. *Pnky*-KO females exhibit decreased PPI at the prepulse intensity, 4dB over the background; \**p* = 0.0202, multiple *t* tests. PPI was not altered in male Pnky KO mice (multiple *t*-tests, *p =* ns for all prepulse intensities). \**p* < 0.05, *p* < 0.01, ns = non-significant, data is represented as mean ± SEM.

Prepulse inhibition (PPI) of ASR is the reduction in the amplitude of the startle response when a weak, non-startling tone – the prepulse – precedes the loud, startling tone. PPI is considered to represent a type of sensorimotor gating, wherein sensory information is “filtered” in the brain before a motor or cognitive response (Gómez-Nieto, Hormigo, and López 2020). To measure PPI, mice are exposed to a series of acoustic startle tones (120 dB) in which some are preceded by weaker prepulse tones of different intensities (4, 15, and 26 bB) above background noise. Startle amplitude is quantified, and for each prepulse intensity, PPI is represented as the percent reduction of the startle amplitude as compared to the baseline (non-prepulse) startle amplitude. In male *Pnky*-KO mice, PPI across the different prepulse intensities was not different from that of their WT controls (**Figure 4B**). In contrast, at the lowest prepulse intensity (4 dB above background) female *Pnky*-KO mice exhibited reduced PPI (**Figure 4B**), indicating that *Pnky* is required for normal PPI in female mice.

### BAC-*Pnky* reduces the acoustic startle response of female *Pnky*-KO mice

BAC-*Pnky* rescues transcriptomic, cellular and neurodevelopmental phenotypes of *Pnky* deletion. However, whether a lncRNA expressed in *trans* can reverse behavioral phenotypes – which are fundamentally more complex than molecular, cellular and even developmental phenotypes – has been unclear. We therefore generated a cohort of *Pnky*-KO (15 females and 18 males) and *Pnky*-KO;BAC-*Pnky* (13 females and 16 males) littermates for behavioral testing during the 5th to 6th month after birth (**Figure 5A**).

**Figure 5.**
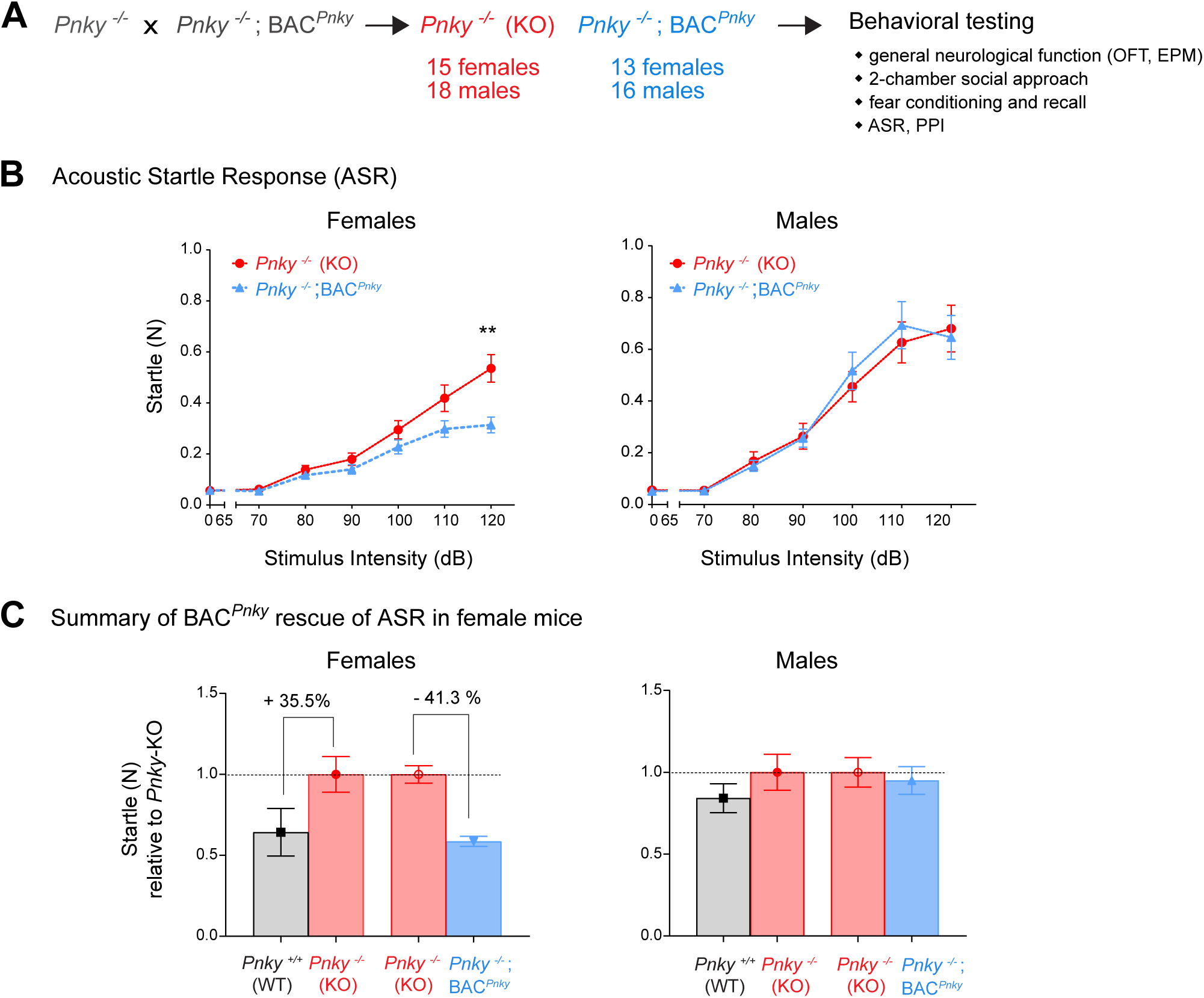
BAC-*Pnky* reduces the acoustic startle response of female *Pnky*-KO mice. **A**) *Pnky*-/-mice were crossed to *Pnky*-/-; BAC-*Pnky* mice to obtain a cohort of *Pnky*-KO (n= 15 females and 13 males) mice and littermate *Pnky*-KO;BAC*-Pnky* ( n= 18 females and 16 males). **B**) *Pnky* KO; BAC-*Pnky* females exhibit decreased startle compared to its KO littermates (rank summary analysis overall effect \**p =* 0.0195 across the whole range of stimuli, followed by Welch’s *t*-tests at 0dB – *p* = ns, and 120dB – \*\**p* = 0.0018; Mann-Whitney test at 70dB – *p* = 0.0907 (ns), 80dB – *p* = ns, 90 dB – *p* = ns, 100 dB – *p* = ns, 110 dB – *p* = ns) while for the males there is no significant difference (rank summary analysis overall effect *p =* ns across the whole range of stimuli). Data is represented as mean ± SEM. **C)** ASR reduction observed with BAC-*Pnky* in *Pnky*-KO female mice (41.3%) is comparable to the amount of ASR increase observed with *Pnky*-KO vs. WT (35.5%) at 120dB.

General neurological function as assessed by OFT was not different between *Pnky*-KO and *Pnky*-KO;BAC-*Pnky* mice in either sex (**Figure 5** **supplement 1A**). Assessment of anxiety and social interactions with the EPM and two-chamber social approach tests, respectively, also did not reveal behavioral differences between *Pnky*-KO and Pnky-KO;BAC-*Pnky* mice (**Figure 5** **supplement 2 A-B**). Thus, BAC-*Pnky* does not produce behavioral changes in *Pnky*-KO mice among behaviors that were not different in *Pnky*-KO mice versus the WT controls.

Male *Pnky*-KO were deficient in the cognitive test of cued fear recall, but BAC-*Pnky* did not change fear conditioning or recall in *Pnky*-KO mice in either male or female mice ( **Figure 5** **supplement 3A-C**). Similarly, while *Pnky*-KO female mice exhibited lower PPI as compared to their WT controls, BAC-*Pnky* did not increase PPI in either sex as compared to the *Pnky*-KO littermates (**Figure 5** **supplement 4A**). These data indicate that BAC-*Pnky* cannot reverse certain behavioral phenotypes related to *Pnky*-KO.

When BAC-*Pnky* was present, female *Pnky*-KO mice exhibited a strong decrease in ASR (**Figure 5B**). In contrast, in male *Pnky*-KO mice, the presence of PAC-*Pnky* did not change the ASR (**Figure 5B**). The amount of ASR reduction (percent change) observed with BAC-*Pnky* in *Pnky*-KO female mice was very similar to the amount of ASR increase observed with *Pnky*-KO vs. WT (**Figure 5C**), suggesting that BAC-*Pnky* produces a nearly complete reversal of the *Pnky*-KO ASR phenotype of female mice.

## DISCUSSION

Animal behavior results from a complex interplay between the genome and the environment. Although many protein coding genes are known to be important to behavior by regulating brain development and/or function, the degree to which lncRNA genes underlie animal behavior is unclear. Relatively few lncRNAs have been studied in mice with genetic methods (*e.g.,* gene disruptions and/or germline transgene insertions), and for only a handful of lncRNAs have the behavioral consequences of such genetic manipulations been evaluated (Oliver et al. 2015; Ip et al. 2016). In this study, we discovered that *Pnky*-KO mice have sex-specific behavioral deficits in fear conditioning and recall, ASR and PPI. Furthermore, the expression of *Pnky* from a BAC transgene reversed the female-specific ASR phenotype of *Pnky*-KO mice, providing molecular-genetic evidence that *Pnky* can underlie animal behavior by functioning in *trans*.

The amplitude of the startle response is an important measure for evaluating affective deficits in patients with specific brain lesions (amygdala and frontal cortex), psychopathic tendencies, and a range of psychiatric conditions including panic disorder, bipolar disorder, and posttraumatic stress syndrome (Angrilli et al. 1996; Poli and Angrilli 2015; Koch 1999; Jovanovic et al. 2009; Giakoumaki et al. 2010). More specifically, increased startle amplitude is associated with greater anxiety levels in young female women (Poli and Angrilli 2015). While female *Pnky*-KO mice had increased ASR amplitude, we did not observe differences in levels of anxiety as assessed by the OFT and EPM. Furthermore, nest building behavior – an indicator of well-being in mice – was not altered in *Pnky*-KO mice of either sex. Thus, the change in ASR in female mice lacking *Pnky* is not obviously related to a general increase in anxiety or change in the sense of well-being.

The ASR in rodents has been found to vary by biological sex, with male rats having increased ASR amplitude as compared to females (Plappert, Rodenbücher, and Pilz 2005; Lehmann, Pryce, and Feldon 1999). In WT mice, we also observed higher ASR amplitude in males versus females. While *Pnky*-KO increased ASR amplitude in females, no difference in ASR was observed in males across a wide range of acoustic intensities. Given that ASR is not modulated by the estrous cycle (Plappert, Rodenbücher, and Pilz 2005), we suggest that this important aspect of biological sex differences does not underlie the sex-specific role of *Pnky* in ASR.

In female *Pnky*-KO mice, BAC-*Pnky* reduced ASR amplitude, which can be interpreted as a reversal of this *Pnky*-KO behavioral phenotype. Because we were unable to conduct the scale of animal husbandry required to generate the necessary numbers of all experimental groups to assess rescue (*i.e., Pnky*-WT, *Pnky*-WT;BAC-*Pnky*; *Pnky*-KO, and *Pnky*-KO;BAC-*Pnky*), we are unable to fully demonstrate “rescue” of the *Pnky*-KO phenotypes. However, we find it compelling that the reduction in ASR amplitude by BAC-*Pnky* was not observed in male mice, indicating that this behavioral effect was specific and not reflective of a more general effect on ASR by the BAC-*Pnky* transgene. To our knowledge, this is the first example of a genetic study demonstrating *trans* function for a lncRNA in animal behavior.

Not all behavioral phenotypes of *Pnky*-KO mice were reversed by BAC-*Pnky*. Neither cued fear recall nor PPI in *Pnky*-KO mice were changed by BAC-*Pnky* in either sex. We acknowledge that it remains possible that *Pnky* functions partly in *cis*. However, another possible explanation could be that *Pnky* expression in *Pnky*-KO;BAC-*Pnky* mice does not fully recapitulate the levels and/or timing of *Pnky* expression in WT mice. *Pnky* is a nuclear-enriched lncRNA transcript, and although BAC-*Pnky* derived transcripts are also nuclear and at present at WT levels of abundance in the *Pnky*-KO;BAC-*Pnky* brain, we have noted transiently lower levels (20% of WT) in cell culture studies after acute *Pnky*-deletion. Thus, we speculate that certain behaviors require more precise regulation of *Pnky* expression for rescue in our assays.

Despite the lack of rescue by BAC-*Pnky*, the sex-specific behavioral deficits in cued fear recall and PPI related to *Pnky*-KO are interesting to consider in the context of studies linking aberrant lncRNA expression and psychiatric diseases. Abnormal fear conditioning and recall is associated with anxiety, depression and many other mental health conditions (Dibbets, van den Broek, and Evers 2015). A reduction in PPI, which is believed to be caused by defects in sensorimotor gating, is considered to be a behavioral biomarker of schizophrenia (Mena et al.

2016; Gómez-Nieto, Hormigo, and López 2020). Interestingly, in a small study of human blood samples, *PNKY* was found to be a strong molecular biomarker of treatment-resistant schizophrenia (Badrlou et al. 2021). Overall, our finding that *Pnky*-KO mice are behaviorally grossly normal but exhibit phenotypes suggestive of affective disorders support the broader notion that lncRNAs in the brain serve to “fine-tune” the development and/or functions that underlie more complex behaviors.

In this study, we focused on behavioral studies and have not associated results with potential anatomic differences or alterations in neural cell function. Based on the literature, much of which is correlative, the underpinnings of the observed functional differences are very interesting but numerous. For instance, differences in ASR could relate to differences in amygdala or orbitofrontal cortex anatomy and/or function, as these areas provide afferent modulation to the brainstem ASR circuits (Koch 1999). *Pnky* is also expressed in the adult mouse brain, and it is therefore possible that this lncRNA modulates adult neuronal function. However, based on our previous findings of abnormalities in the postnatal cortex of *Pnky*-KO mice (Andersen et al. 2019), we currently hypothesize that specific anatomic differences in cell type and/or connectivity underlie the observed behavioral phenotypes.

Functional studies linking lncRNAs with animal behaviors comprise an emerging field of research that is important to our understanding of a wide range of neurological diseases including cognitive and psychiatric disorders. Because the breadth of potential mechanisms of lncRNAs is fundamentally wider (*cis* and *trans*) than most protein coding or microRNA genes, the strategies used to study lncRNA function may need to be tailored to each lncRNA (Bassett et al. 2014; Kopp and Mendell 2018; Mattick et al. 2023). The study of animal behavior is also experimentally complex, with results being sensitive to a large number of potential variables, and genetic tools are a powerful means to reducing the variability of lncRNA perturbation. Moving forward, additional *in vivo* studies of lncRNA function in animal behavior will become a crucial foundation for understanding how specific lncRNAs can underlie important disorders of the human mind.

## Acknowledgements

We thank members of the Lim lab for helpful discussions and Sandra Chang for work with animal husbandry and administrative expertise. We thank Iris Lo, Julia Holtzman, and Jessica Speckart for contributions to the behavioral analyses within the Gladstone Institutes Behavioral Core Facility. This work was supported by NIH award 1R01NS124881 and Veterans Affairs 5I01 BX000252 to D.A.L. Figures created with

## Author Contributions

R.E.A. and D.A.L. conceptualized the study and designed experiments with guidance from J.S. P.S., R.E.A., J.S. and D.A.L. analyzed data, and P.S. and D.A.L. prepared the figures. S.J.H. and E.G. performed and analyzed experiments. P.S. and D.A.L. wrote the manuscript. All authors reviewed and edited the manuscript.

## Materials and Methods Animals

All C57BL/6J mice were group-housed and maintained in the University of California, San Francisco Laboratory Animal Resource Center under protocols approved by the Institutional Animal Care and Use Committee. All relevant ethical regulations were followed. Mice of both sexes were used for all experiments and were analyzed at ages 5-6 months. All test results were analyzed relative to littermates. Animals were ear-tagged, and the experimenter of the behavioral tests was blinded to the genotype of the animals.

## Open field Test

Open field activity is measured in a clear acrylic chamber (41 x 41 x 30 cm) with two 16 x 16 photobeam arrays that automatically detect horizontal and vertical movements, also known as the Flex-Field/Open Field Photobeam Activity System (San Diego Instruments, San Diego, CA). The acrylic chambers are located inside larger sound and light attenuating shells so that their spontaneous locomotor activity is not affected by external stimuli. Mice are allowed to habituate in the testing room under normal light for 60 min before testing. During testing, mice are placed in the center of the activity chamber and allowed to freely explore for 15 minutes.

## Elevated Plus Maze Test

The elevated plus maze consists of two open arms (without walls, 15” long X 2” wide) and two closed arms (with walls 6.5” tall), the intersection of the arms is 2” x 2” wide, and the entire maze is elevated 30.5” above the ground (Hamilton-Kinder, Poway, CA). Mice are first allowed to habituate in the testing room under dim light for 1 hr prior to the start of testing. During testing, mice are placed in the maze at the intersection of the open and closed arms and allowed to freely explore the maze for 10 min. The maze is cleaned with 70% alcohol between testing of each mouse. Main dependent measures are as follows 1) % Open arm time 2) Open arm/total distance 3) Closed arm distance 4) Total distance 5) Open arm entries 6) Closed arm entries 7) % Open arm time x minutes.

## Rotarod test

The training part of the test is used to introduce the mouse to the rotarod apparatus (Med Associates Inc., Vermont, USA) which is equipped with infrared beams that automatically detect when the mouse has fallen off the rotating rod. Each day of testing, the mice are allowed 1hr to habituate to the procedure room before testing begins. On the first day, up to five mice of the same sex are simultaneously placed on the rotarod apparatus, and then it rotates at the constant speed of 16 RPM. The trial ends when the mouse falls off the rod or when 5 minutes has elapsed. The mice are tested on 3 individual trials with an inter-trial interval between 15-20 minutes.

The testing phase assesses their general motor learning ability or increase in performance over trials. On the second and third day of testing, five mice of the same sex are simultaneously placed on the rotarod apparatus with the rod rotating at an accelerated speed, from 4 RPM to 40 RPM. The rotation speed increases by 7.2 RPM every minute. The trial ends when the mouse falls off the rod or when 5 minutes has elapsed. The inter-trial interval is between 15-20 minutes. For the two testing days, the animals get a total of 6 trials per day separated into two 3-trial sessions with inter-session interval of ∼3 hours.

## Nesting

A standard mouse cage (10” x 7” x 6.5”) is filled with ∼2cm of paper chip bedding and a single nestlet (5cm square of pressed cotton batting) is placed in the center of the cage. Each mouse is single housed for the 24hr testing period and the quality of their nest is scored at 2, 6, and 24 hrs after introduction of the nestlet. Nests are assigned a score from 0 to 7 at each time point. Scoring: 0 = nestlet untouched, 1 = less than 10% of the nestlet is shredded, 2 = 10-50% of the nestlet is shredded but there is no shape to the nest, 3 = 10-50% of the nestlet is shredded and there is shape to the nest, 4 = 50-90% of the nestlet is shredded but there is no shape to the nest, 5 = 50-90% of the nestlet is shredded and there is shape to the nest, 6 = Over 90% of the nestlet is shredded but the nest is flat, 7 = Over 90% of the nestlet is shredded and the nest has walls that are as tall as the mouse on at least 50% of its sides.

## 2-trial Social Approach Test

Mice are brought into testing room and given one hour to acclimate to the room prior to testing. The social arenas are constructed from white acrylic. Dividers are clear acrylic with arch-shaped entrances at the center. For 2-chamber, arenas are divided into two 30W x 40D x 23H cm chambers. The social arenas are housed inside sound attenuating shells measuring 80W x 67D x 64H cm and equipped with two 0.5-amp lights and an exhaust fan. Social & non-Social enclosures are circular wire cups 10cm in diameter and 13cm tall. The spacing of the wire allows for olfactory and visual access between the stimulus and experimental mice. All stages of the experiment are videotaped from above using cameras mounted on the top of the sound attenuating shells. Mice are allowed 1hr to acclimate to the procedure room before testing begins. The habituation phase and testing phase occurs back-to-back. During habituation the mouse is allowed to explore the entire arena with empty enclosures for ten minutes. During social approach trial, the mouse is allowed to freely explore the entire arena for another 10 minutes, but the enclosures now contain an inanimate mouse on one side and a live social stimulus mouse (non-transgenic, age and strain matched, non-cage/littermate) on opposite side. The experimental mouse is blocked in the nonsocial chamber before introducing the stimulus mouse and inanimate mouse into their respective enclosures. The location of the stimulus mouse is counterbalanced across genotypes. Videos are analyzed using CleverSys TopScan.

The time and bouts spent in each chamber, in the proximity zone (5 cm area perimeter around enclosure), and “sniffing” (1.5 cm area perimeter around enclosure) is recorded. Difference scores (Social minus Non-Social) for bouts and time are calculated for the chamber side and interaction/proximity zones. Ratio scores (Social divided by Non-Social) for bouts and time are calculated for the chamber side and interaction/proximity zones.

## Object-context congruence test

Mice are moved into the testing room for 1 hr to acclimate under normal lighting conditions. Experiment is carried out inside sound attenuating shells equipped with two 5W houselights, and an overhead video camera. Mice are randomly distributed into two groups: Context A first or Context B first for Trial 1, and reverse order for Trial 2. For the test phase, all the animals were run in the context that they had seen most recently (training trial 2) but with an incongruent object from the other context. Context A Chamber: plain white, Objects: white rectangular blocks, Cleaning Agent: 70% EtOH. Context B Chamber: checkered walls, Objects: 50mL Erlenmeyer flasks with tented tops (filled with colored paper), Cleaning Agent: 1% Acetic Acid.

Each trial is videotaped from above and videos are analyzed with post-acquisition software, CleverSys TopScan, to measure the time spent sniffing the objects within each Context (<1.5 cm distance around objects.). The objects are either cleaned with 70% EtOH or 1% Acetic Acid (dependent on Context) in between each mouse. Mice are tested in 1 day using three 10-minute trials: 1) Trial 1 (Object-Context A or B pairing) for 10 min exploration then home cage for 30 minutes. 2) Trial 2 (Object-Context B or A pairing) for 10 min exploration then home cage for 4 hrs. 3) Object-Context Congruence Test in the most recent context with one congruent and one incongruent 10 min exploration then home cage.

## Cued fear conditioning and recall

Testing is done in the fear conditioning chamber 9.5” L x 12” W x 8.5” H (Med Associates) that sits inside a sound attenuating shell 25” L x 29.5’ W x 14” H. The Med Associates VideoFreeze program is used for tracking. Mice are not habituated to the testing room on the day of testing and are brought into the testing room immediately before the trial begins.

Training (Day 1): Mice are placed using nitrile gloves in the fear conditioning apparatus with light set at 3, fan on, tray is sprayed with Windex, and the floor is an even-barred metal grid.

Test starts with a 5 min baseline period to measure baseline freezing activity. Then four-30 second 80dB tones that co-terminate with a 2-second, 0.45mA footshock are presented, separated by 120-second inter-cue interval (ICI) during which freezing is monitored. A 120 second ICI follows the last footshock. Fear context test (Day 2): The context test takes place 24 hours after Training. Setup is the exact same as training. Mice are placed in the fear conditioning apparatus for 10 minutes. No cues or shocks are presented. Freezing is monitored. Cued fear recall Test (Day 3): takes place 24 hours after the context test. The context is altered:

A black insert covers the metal grid floor, and a black A frame is added. Chamber light set at 1, fan is off, and room lights are dimmed. Mice are placed with latex gloves into a different chamber from the training and context test phases for 15 minutes. After a 5 min context generalization period, four tones are delivered as described in Training. No shock is presented.

## Acoustic Startle Response

Testing is performed in a small, isolated chamber inside a sound attenuating cubicle, free from external movement and noise (Hamilton-Kinder, Poway, CA). On the day of testing, an individual cage of group-housed mice or two single-housed mouse cages is/are transferred to the ante room just outside and prior to testing. The startle sensor plate is calibrated at the start of each testing day. The mice are placed into the restraining chamber and given 5 minutes to acclimate inside that restraining chamber before stimulus presentations begin. The restraining chamber is cleaned with 70% ethanol in between individual mice.

Habituation: Mice are given 5 minutes to acclimate to the restraining chamber and 64 dB background noise before acoustic stimulus testing begins. After 5 minutes, mice are exposed to a series of acoustic pulses at varying intensities for approximately 15 minutes at random intervals. Trials with no auditory stimulus are also included to provide a measure of baseline startle activity. The test is comprised of a total of 70 trials that are randomly selected from the following list: 10 trials each of 40ms @120dB, 40ms @110dB, 40ms @100dB, 40ms @90dB, 40ms @80dB, 40ms @70dB and no stimulus trials. The interstimulus interval (ISI) is variable with a mean of 15 seconds, and a range of 8-22 seconds. Average and Maximum amplitudes (N) of pulses are measured for each mouse.

## Pre-pulse Inhibition

Testing is performed in a small, isolated restraining chamber placed inside a sound attenuating cubicle, free from external movement and noise (Hamilton-Kinder, Poway, CA). The startle sensor plate is calibrated at the start of each testing day. Habituation: Mice are given 5 minutes to acclimate to the restraining chamber and 64 dB background noise before acoustic stimulus testing begins. After 5 minutes, mice are exposed to a series of acoustic startle stimuli for 20 minutes in which some stimuli are preceded by a weaker acoustic stimulus (pre-pulse) at random intervals. Trials with no auditory stimulus are also included to provide a measure of baseline activity. The test is comprised of a total of 80 trials randomly selected from the following list: 24 trials of 40ms@120dB, and 14 trials each of the 4dB pp 40ms @120dB, 15dB pp 40ms @120dB, 26dB pp 40ms@120dB, and no stimulus trials. The interstimulus interval (ISI) is variable with a mean of 15 seconds, and a range of 8-22 seconds. Average and Maximum amplitude of startle responses to each acoustic stimulus (measured in “N” calibrated units), as well as those with preceding pre-pulses, are recorded for each animal.

## Figure Legends

**Figure 1 Supplement 1: A)** Ambulatory movement in open field test is comparable in both sexes and both genotypes (Welch’s *t* test; females *p* = 0.0933 (ns), males *p* = ns). **B)** Rearing behavior shows no significant difference between *Pnky*-WT and *Pnky*-KO animals (Mann-Whitney tests, *p* = ns). Data is represented as mean ± SEM.

**Figure 2 Supplement 1 A)** *Pnky*-WT and *Pnky*-KO mice have comparable center to total movement ratio in the open field test (Welch’s *t*-tests, *p* = ns). **B)** In the elevated plus maze test, females and male animals show no significant genotype-specific difference in number of entries in the closed arm (Welch’s *t*-tests, *p* = ns) and distance travelled in the closed arms (Welch’s *t*-tests, *p* = ns). Data is represented as mean ± SEM.

**Figure 3 Supplement 1 A)** *Pnky*-KO and WT mice did not exhibit significant differences in the two training trials of the object context congruence task (paired *t*-tests, *p* = ns). **B)** Female and male mice of both genotypes show increased preference (% interaction time in seconds) for the incongruent object whereas the difference in interaction time across genotypes is non-significant. Paired *t* test \*\**p* = 0.0025 for *Pnky*-WT females congruent vs incongruent. Paired *t* test \*\**p* = 0.0051 for *Pnky*-KO females congruent vs incongruent. Paired *t* test \**p*= 0.0109 for *Pnky*-WT males and \*\*\*\**p* <0.0001 for *Pnky*-KO males, congruent vs incongruent. Percent incongruent interaction time was not different between the *Pnky-*WT and *Pnky*-KO of either sex (Welch’s *t*-tests, *p* = ns), \**p* < 0.05, \*\**p* < 0.01, \*\*\*\**p* <0.0001, ns = non-significant, data is represented as mean ± SEM.

**Figure 3 Supplement 2 A)** Cued Fear conditioning test starts with a 5 min baseline period to measure baseline freezing activity. Then four-30 second 80dB tones that co-terminate with a 2-second, 0.45mA foot shock are presented. Freezing behavior in *Pnky*-KO mice was comparable to its WT controls (rank summary analysis for male and female baseline; multiple *t*-tests during the cue presentations for female and male mice, *p* = ns; multiple *t*-tests for male and female ICI, *p* = ns). **B)** In cued fear recall test After a 5 min context generalization period, four tones are delivered as described in fear conditioning. No shock is presented. *Pnky*-WT and *Pnky*-KO females exhibit no significant difference in freezing behavior (linear mixed effects analysis for generalization; multiple *t*-tests during the cue presentations *p* = ns; linear mixed analysis for ICI 1, 3, and 4, *p* = ns; rank summary analysis for ICI 2*, p* = ns). *Pnky*-KO male mice show decreased freezing compared to the WT control in the 5-minute generalization period (rank summary analysis, \**p* = 0.0191), during the cue presentations (multiple t-tests, \**p* = 0.0301) and during the ICI (linear mixed effects analysis revealed significant differences for ICI 1 and 4, \**p* = 0.0237 and \**p* = 0.0478, respectively, but not for ICI 2 and 3, p = ns)). \**p* < 0.05, ns= non-significant, data is represented as mean ± SEM.

**Figure 5 Supplement 1 A)** In the OFT, no significant differences between females and males on *Pnky*-KO and *Pnky*-KO; BAC-*Pnky* groups in total movement (repeated measures two-way ANOVA, *p* = ns) and ratio of center to total movement (Welch’s *t*-tests, *p =* ns). Data is represented as mean ± SEM.

**Figure 5 Supplement 2 A)** In the EPM, the ratio of open arm distance to total distance (females, Mann-Whitney, *p* = ns; males, Welch’s *t*-test, *p =* ns), number of open arm entries (Welch’s *t*-tests, *p =* ns), number of closed arm entries (Welch’s *t-*tests, *p =* ns), and closed arm distance (Welch’s *t*-tests, *p =* ns) are comparable between the *Pnky*-KO and *Pnky*-KO; BAC-*Pnky* groups. **B)** Female and male mice of both genotypes show increased preference (interaction time in seconds) for the social side in the 2-trial social approach test whereas the difference in interaction time across genotypes is non-significant. Paired *t* test \*\*\*\**p*= 0.0001 for social vs inanimate side for both sexes for both *Pnky*-KO and *Pnky*-KO; BAC-*Pnky* groups. Welch’s *t*-test *p* = ns for *Pnky*-KO vs *Pnky*-KO; BAC-*Pnky* social interaction time. \*\*\*\**p* <0.0001, ns = non-significant, data is represented as mean ± SEM.

**Figure 5 Supplement 3** Females and males of *Pnky*-KO and *Pnky*-KO; BAC-*Pnky* groups show no significant difference in **A)** cued Fear conditioning (multiple *t*-tests during baseline, the cue presentations, and the ICI for the female and male mice, *p =* ns) **B)** fear context recall (Welch’s *t*-test, *p =* ns), and **C)** cued fear recall tests (multiple *t*-tests during the context generalization, cue presentations, and ICI for female and male mice, *p* = ns)

**Figure 5 Supplement 4 A)** *Pnky*-KO and *Pnky*-KO; BAC-*Pnky* animals exhibit no significant changes in the startle inhibition at prepulse intensities, 4dB, 15dB and 25dB over the background (multiple *t*-tests for each sex, *p* = ns).

