## SupplementaryFigures for "Sex-specific role for the long noncoding RNA *Pnky* in mouse behavior"

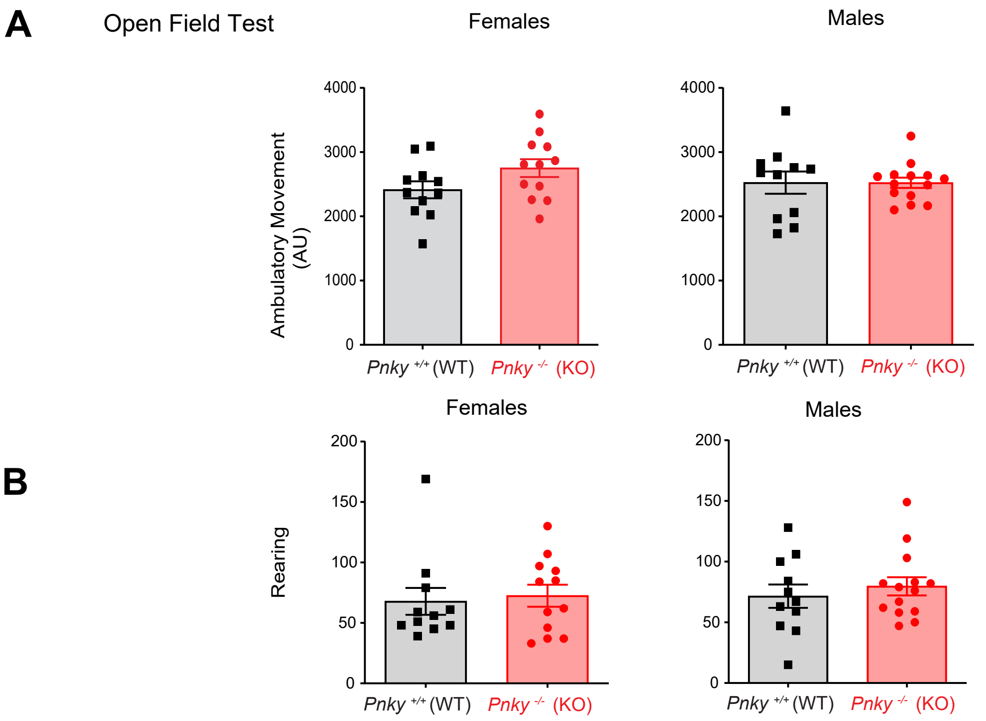


**Figure 1 Supplement 1: A)** Ambulatory movement in open field test is comparable in both sexes and both genotypes (Welch's *t* test; females $p = 0.0933$ (ns), males $p$ = ns). **B)** Rearing behavior shows no significant difference between *Pnky*-WT and *Pnky*-KO animals (Mann-Whitney tests, $p$ = ns). Data is represented as mean ± SEM.

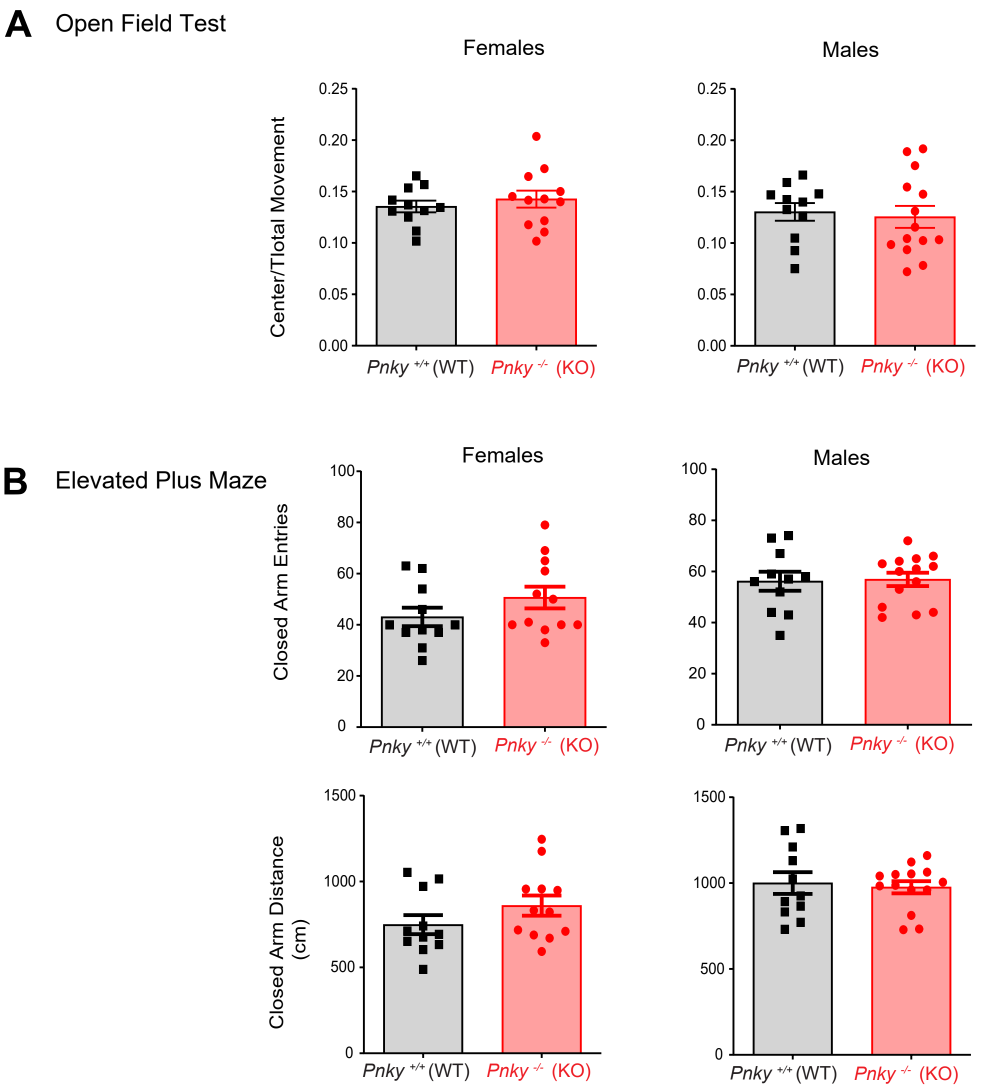


**Figure 2 Supplement 1 A)** *Pnky*-WT and *Pnky*-KO mice have comparable center to total movement ratio in the open field test (Welch's *t*-tests, $p$ = ns). **B) I**n the elevated plus maze test, females and male animals show no significant genotype-specific difference in number of entries in the closed arm (Welch's *t*-tests, $p$ = ns) and distance travelled in the closed arms (Welch's *t*-tests, $p$ = ns). Data is represented as mean ± SEM.

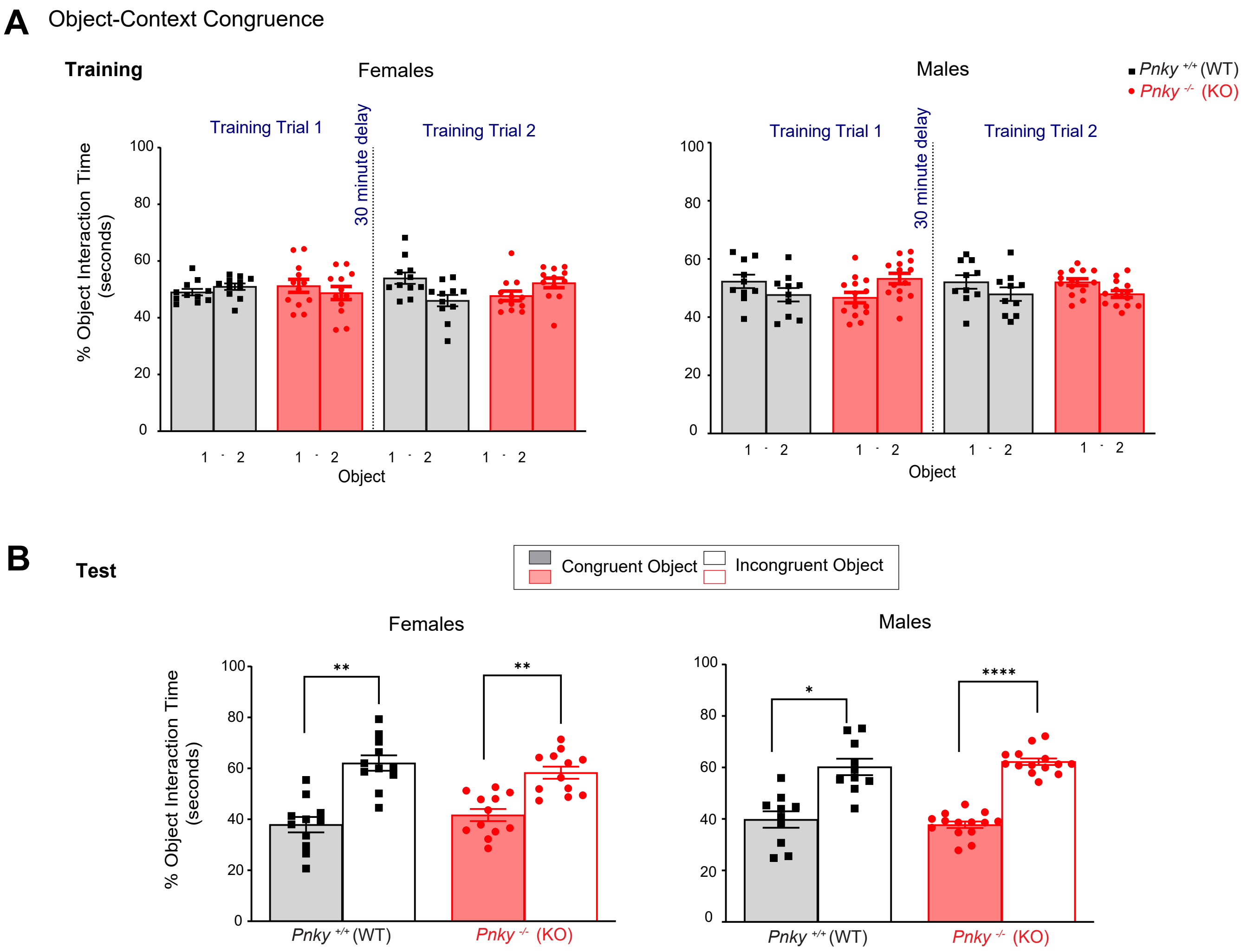


**Figure 3 Supplement 1 A)** *Pnky*-KO and WT mice did not exhibit significant differences in the two training trials of the object context congruence task (paired *t*-tests, $p$ = ns). **B)** Female and male mice of both genotypes show increased preference (% interaction time in seconds) for the incongruent object whereas the difference in interaction time across genotypes is non-significant. Paired *t* test $^{**}p = 0.0025$ for *Pnky*-WT females congruent vs incongruent. Paired *t* test $^{**}p = 0.0051$ for *Pnky*- KO females congruent vs incongruent. Paired *t* test $^{*}p= 0.0109$ for *Pnky*-WT males and $^{****}p <0.0001$ for *Pnky*-KO males, congruent vs incongruent. Percent incongruent interaction time was not different between the *Pnky*-WT and *Pnky*-KO of either sex (Welch's *t*-tests, $p$ = ns), $^{*}p < 0.05$, $^{**}p < 0.01$, $^{****}p <0.0001$, ns = non-significant, data is represented as mean ± SEM.

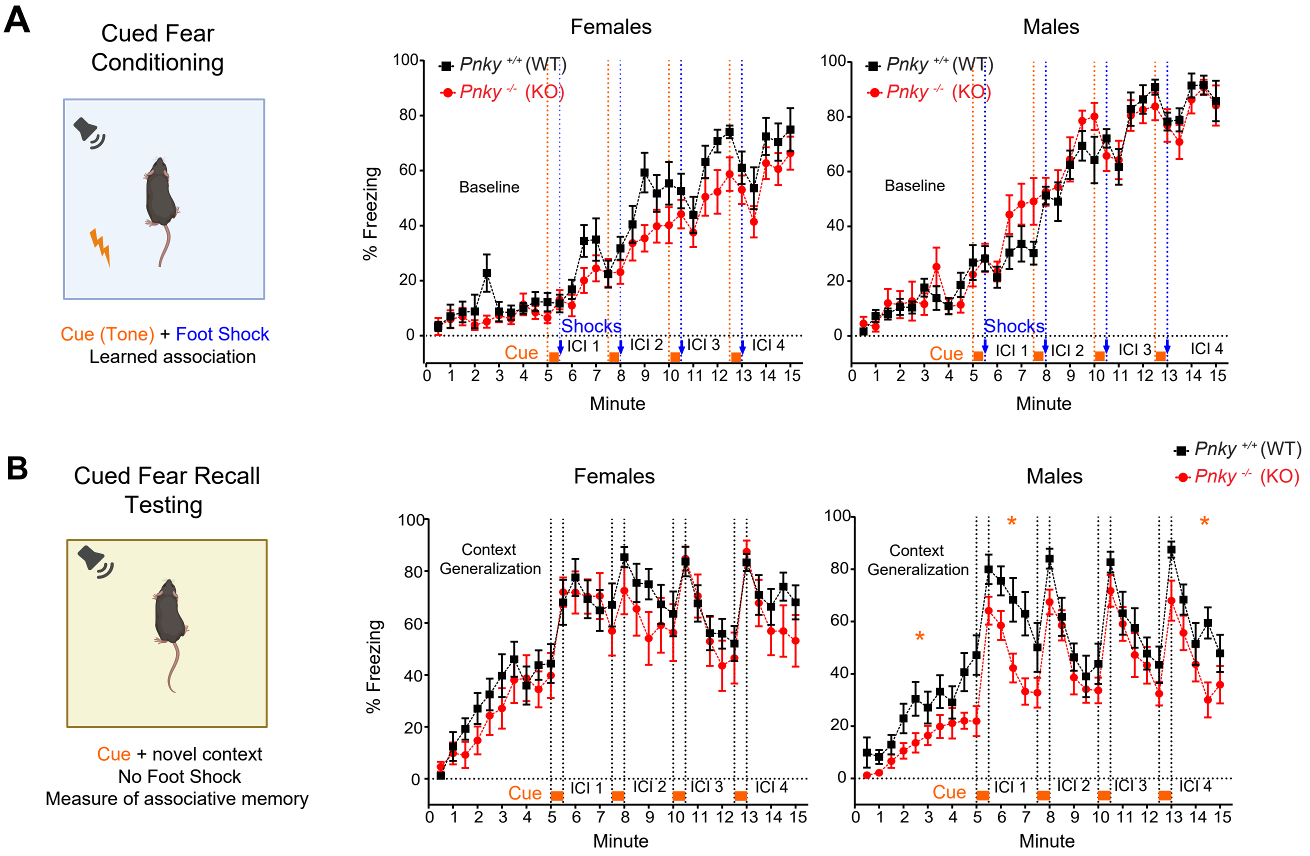


**Figure 3 Supplement 2 A) Cued fear conditioning test** starts with a 5 min baseline period to measure baseline freezing activity. Then four-30 second 80dB tones that co-terminate with a 2-second, 0.45mA footshock are presented,separated by 120-second inter-cue interval (ICI) during which freezing is monitored. A 120 second ICI follows the last footshock. Freezing behavior in *Pnky*-KO mice was comparable to its WT controls (rank summary analysis for male and female baseline; multiple *t*-tests during the cue presentations for female and male mice, $p$ = ns; multiple *t*-tests for male and female ICI, $p$ = ns). **B)** In the **cued fear recall test**, after a 5 min context generalization period, four tones are delivered as described in conditioning phase. No shock is presented. *Pnky*-WT and *Pnky*-KO females exhibit no significant difference in freezing behavior (linear mixed effects analysis for generalization; multiple *t*-tests during the cue presentations $p$ = ns; linear mixed analysis for ICI 1, 3, and 4, $p$ = ns; rank summary analysis for ICI 2, $p$ = ns). *Pnky*-KO male mice show decreased freezing compared to the WT control in the 5-minute generalization period (rank summary analysis, *$p = 0.0191$), during the cue presentations (multiple *t*-tests, *$p = 0.0301$) and during the ICI (linear mixed effects analysis revealed significant differences for ICI 1 and 4, *$p = 0.0237$ and *$p = 0.0478$, respectively, but not for ICI 2 and 3, p = ns)). *$p < 0.05$, ns= non-significant, data is represented as mean ± SEM.

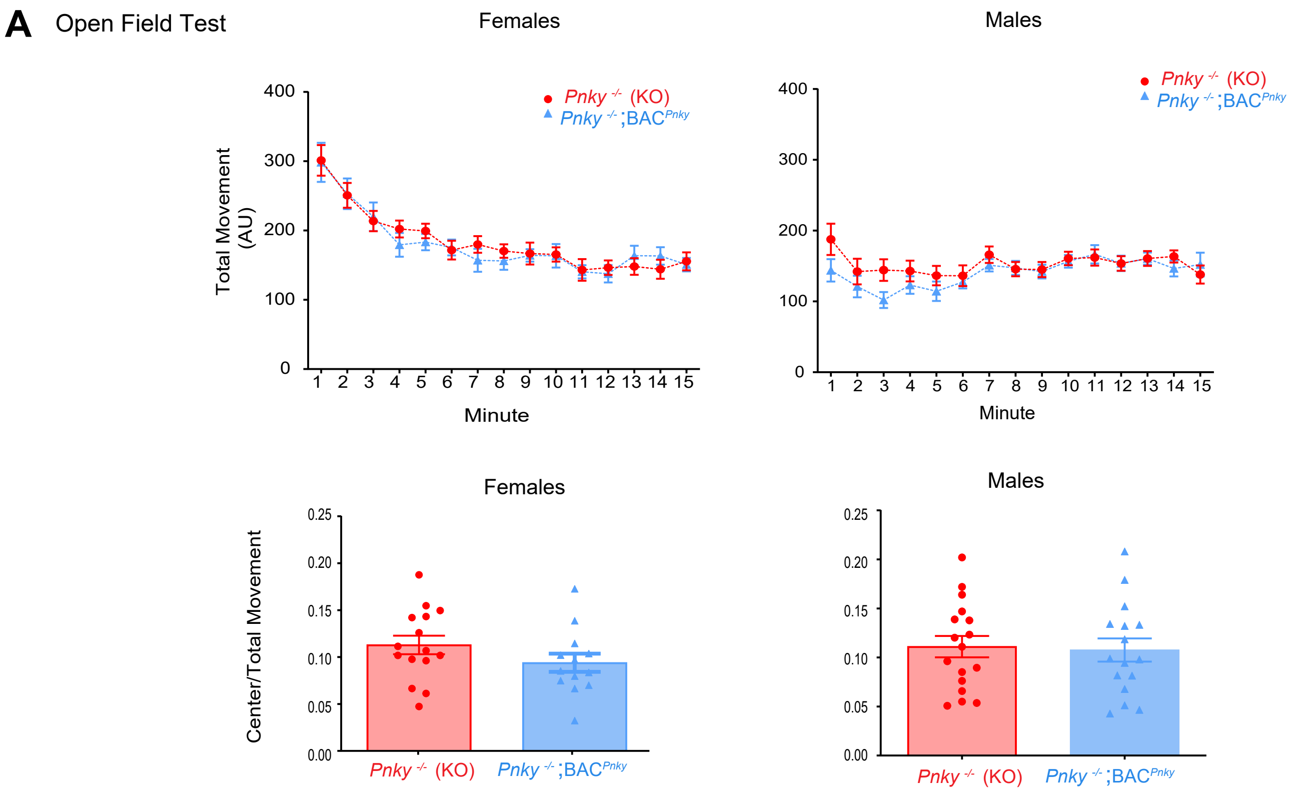


**Figure 5 Supplement 1 A)** In the OFT, no significant differences between females and males on *Pnky*-KO and *Pnky*-KO; BAC-*Pnky* groups in total movement (repeated measures two-way ANOVA, $p$ = ns) and ratio of center to total movement (Welch's $t$-tests, $p$ = ns). Data is represented as mean ± SEM.

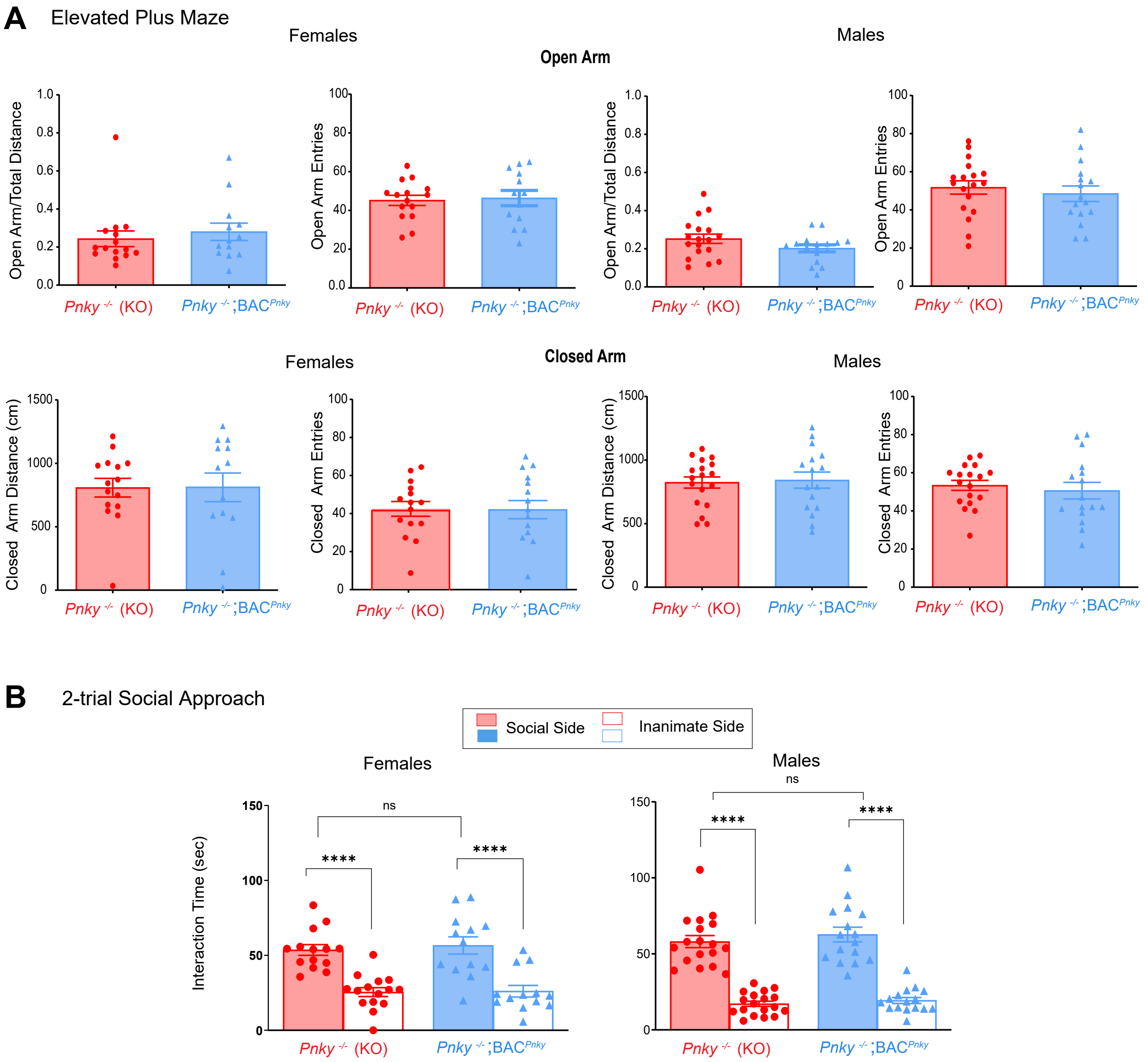


**Figure 5 Supplement 2 A)** In the EPM, the ratio of open arm distance to total distance (females, Mann-Whitney, *p* = ns; males, Welch's *t*-test, *p* = ns), number of open arm entries (Welch's *t*-tests, *p* = ns), number of closed arm entries (Welch's *t*-tests, *p* = ns), and closed arm distance (Welch's *t*-tests, *p* = ns) are comparable between the *Pnky*-KO and *Pnky*-KO; BAC-*Pnky* groups. **B)** Female and male mice of both genotypes show increased preference (interaction time in seconds) for the social side in the 2-trial social approach test whereas the difference in interaction time across genotypes is non-significant. Paired *t* test ****$p$= 0.0001 for social vs inanimate side for both sexes for both *Pnky*-KO and *Pnky*-KO; BAC-*Pnky* groups. Welch's *t*-test *p* = ns for *Pnky*-KO vs *Pnky*-KO; BAC-*Pnky* social interaction time. ****$p$ <0.0001, ns = non-significant, data is represented as mean ± SEM.

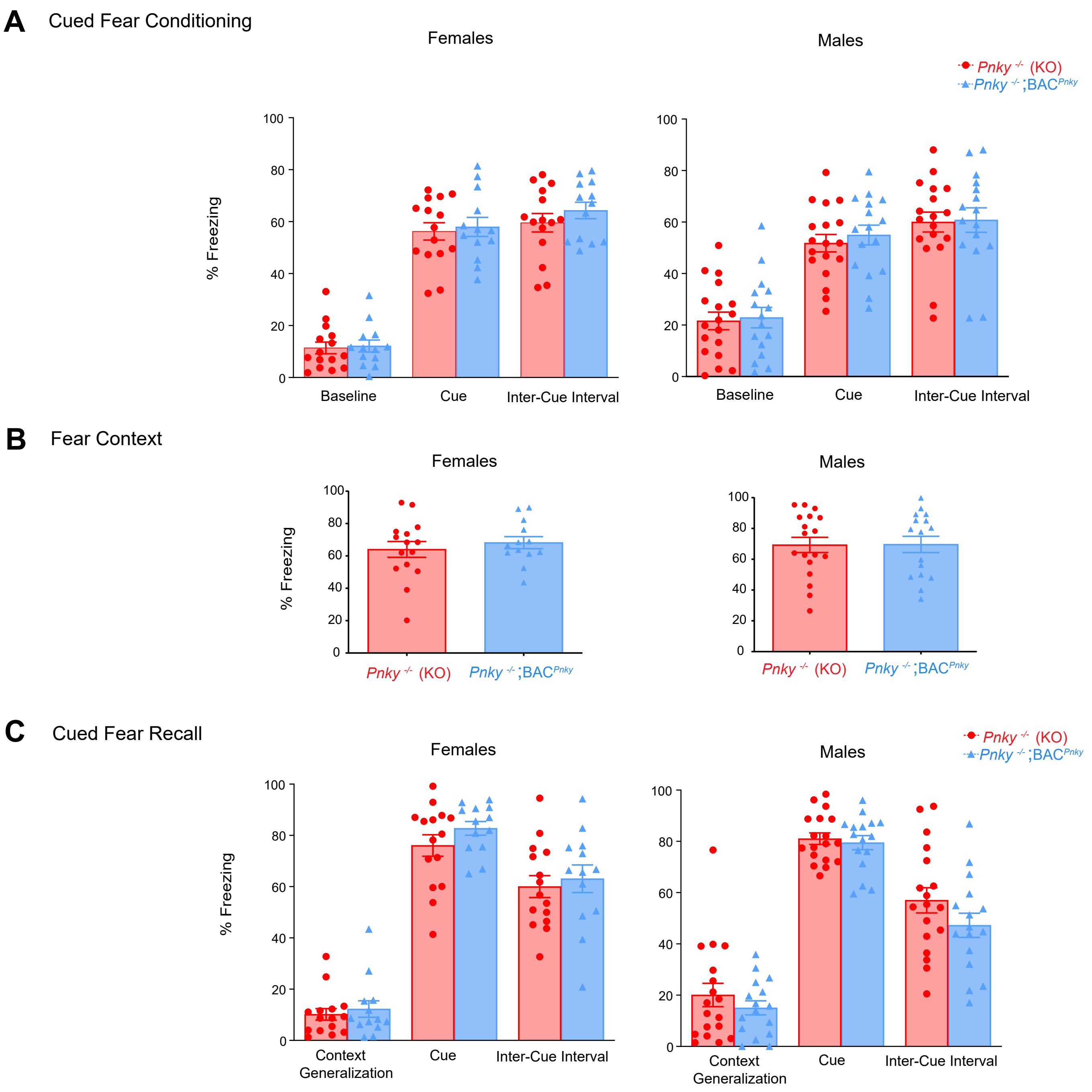


**Figure 5 Supplement 3** Females and males of *Pnky*-KO and *Pnky*-KO; BAC-*Pnky* groups show no significant difference in **A)** cued Fear conditioning (multiple *t*-tests during baseline, the cue presentations, and the ICI for the female and male mice, $p$ = ns) **B)** fear context recall (Welch's *t*-test, $p$ = ns), and **C)** cued fear recall tests (multiple *t*-tests during the context generalization, cue presentations, and ICI for female and male mice, $p$ = ns)

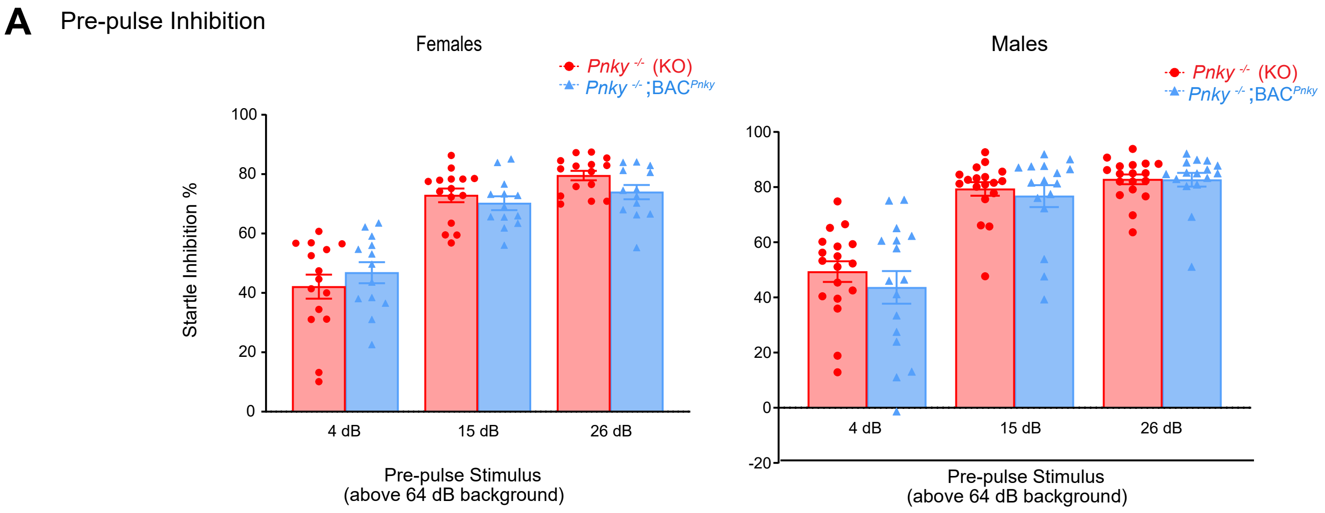


**Figure 5 Supplement 4 A)** *Pnky*-KO and *Pnky*-KO; BAC-*Pnky* animals exhibit no significant changes in the startle inhibition at prepulse intensities, 4dB, 15dB and 25dB over the background (multiple *t*-tests for each sex, $p$ = ns)**.**

.
